# *In vitro* assembly of the Rous Sarcoma Virus capsid protein into hexamer tubes under physiological conditions

**DOI:** 10.1101/098400

**Authors:** Soumeya A. Jaballah, Graham D. Bailey, Ambroise Desfosses, Jaekyung Hyun, Alok K. Mitra, Richard L. Kingston

## Abstract

During a proteolytically-driven maturation process, the ortho-retroviral capsid protein (CA) assembles to form the convex shell that surrounds the viral genome. In some orthoretroviruses, including Rous Sarcoma Virus (RSV), CA carries a short and hydrophobic spacer peptide (SP) at its C-terminus early in the maturation process, which is progressively removed as maturation proceeds. In this work, we show that RSV CA assembles *in vitro* at physiological temperatures, forming hexamer tubes that effectively model the mature capsid surface. Tube assembly is strongly influenced by electrostatic effects, and is a nucleated process that remains thermodynamically favored at lower temperatures, but is effectively arrested by the large Gibbs energy barrier associated with nucleation. RSV CA tubes are multi-layered, being formed by nested and concentric tubes of capsid hexamers. However the spacer peptide acts as a layering determinant during tube assembly. If only a minor fraction of CA-SP is present, multi-layered tube formation is blocked, and single-layered tubes predominate. This likely prevents formation of biologically aberrant multi-layered capsids in the virion. The generation of single-layered hexamer tubes facilitated 3D helical image reconstruction from cryo-electron microscopy data, revealing the basic tube architecture.

## INTRODUCTION

The basic features of ortho-retroviral particle assembly and maturation are well established ^1–5^. Particle formation is driven by the Gag polyprotein - a flexible multi-domain molecule ^6-8^ which packs radially into the assembling virion. Sometime after assembly and budding, cleavage of Gag by the viral protease liberates the major structural proteins of the infectious virus: matrix (MA), capsid (CA), and nucleocapsid (NC), accompanied by a profound change in the internal organization of the viral particle. The defining event in this maturation process is formation of the mature capsid shell, which surrounds the genomic RNA and replicative machinery of the virus.

The mature capsid is constructed predominantly of CA hexamers, with occasional pentamers allowing closure of the capsid. The pentameric and hexameric building blocks of the capsid have been structurally characterized for a number of ortho-retroviruses ^9–17^, including RSV ^18–20^, with the analysis now extending to atomic resolution in several cases ^9–11,14,16^. The pathway of mature capsid formation remains a matter of contention. Most studies support a reconstructive mechanism ^21,22^, involving full disassembly and reassembly of the capsid. However some have proposed a displacive mechanism ^23,24^ in which CA does not freely diffuse during mature capsid formation. *In vitro* assembly studies on purified CA (see ^25^ for review) have certainly established the feasibility of *de novo* capsid assembly, and this study was carried out under assumption of a reconstructive mechanism for mature capsid formation. It addresses some of the open questions that remain regarding the regulation of the assembly process, the induction of curvature into the growing capsid shell, and the overall stability of the assembled capsid.

Irrespective of the mechanism, proteolytic processing of the Gag polyprotein initiates mature capsid formation. Liberation of the N-terminus of CA enables formation of a β-hairpin structure ^26,27^, which appears critical in facilitating assembly of the hexameric and pentameric building blocks of the mature capsid ^28–33^. In this fashion, it has been described as a proteolytic trigger for mature capsid assembly ^34^. In contrast, proteolytic processing at the C-terminus of CA has generally been considered to be a more neutral event, with few specific consequences for mature particle formation. In alpharetroviruses like RSV, and lentiviruses like HIV-1, CA and NC are separated by a short hydrophobic spacer peptide. The proteolytic processing events at the C-terminus of CA are kinetically slow, and are amongst the last to occur during maturation (see ^35,36^ for review). The result, for both RSV and HIV, is that CA species carrying the spacer peptide at their C-terminus (Fig 1) are significantly populated early in the maturation process. The spacer peptide is subsequently slowly trimmed by the viral protease, sometimes being cut internally during this operation.

**Fig 1.**
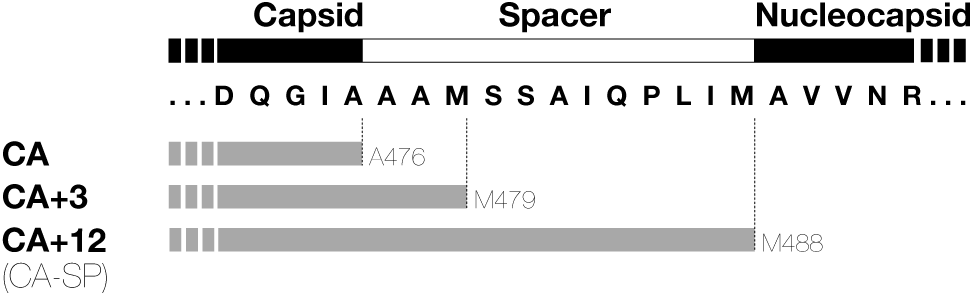
RSV CA variants generated through proteolytic cleavage of the Gag polyprotein. The figure shows the amino acid sequence of the RSV Gag polyprotein (strain Prague C) at the capsid-nucleocapsid junction with the major proteolytic cleavage sites indicated. During maturation, initial cleavage at M488 leads to the transient buildup of the species denoted CA+12 (CA-SP) ^92^. Subsequent proteolytic processing of the spacer peptide yields the species denoted CA and CA+3, which are found in approximately 2:1 ratio in mature RSV virions ^91^.

The spacer peptide has a central role in the oligomerization of Gag, and the stabilization of the immature particle. For RSV, biophysical experiments on the spacer peptide, with short flanking sequences from both CA and NC appended, suggest that this region can self-assemble into a hexameric helical bundle ^37^, consistent with the structure visualized using electron microscopy ^38,39^. Similar results were obtained for the HIV-1 spacer peptide ^40^ and the structure of the hexamer, as it exists in the immature HIV Gag lattice ^41^, was recently determined at atomic resolution ^42,43^. However structure formation by the spacer peptide is clearly context-dependent. For both RSV and HIV-1, studies on unassembled CA and Gag variants have consistently shown that the spacer peptide and immediately neighboring sequences are dynamic, and lack persistent structure ^27,44–48^. The spacer peptide is also disordered within *in vitro* assembled structures that model the mature capsid ^22,49^. Hence the spacer peptide has often been considered irrelevant to mature capsid assembly. Despite this, appending the spacer peptide to RSV CA has been shown to have functional consequences, strongly promoting capsid assembly in *in vitro* assays ^50^.

In this study we demonstrate that incubation of purified RSV CA under physiological conditions causes it to assemble into multi-layered tubes built from capsid hexamers, which effectively model the surface of the mature capsid. We characterize some basic aspects of the tube assembly process, and show that the spacer peptide acts as a layering determinant during tube assembly, effectively blocking the multi-layering which otherwise completely predominates. Finally we report the 3D reconstruction of the single-layered capsid tubes, establishing their basic architecture.

## RESULTS

### Near-physiological temperatures promote assembly of RSV CA into Tubes and Core-Like-Particles

RSV CA was purified to homogeneity, and concentrated to 600-700 μM in a neutral pH buffer containing physiological salt concentrations (150 mM NaCl). Under these conditions the protein is monomeric ^19,27^ and can be stored for extended periods at 4 °C without measurable assembly. The monomeric state of the protein was verified using dynamic light scattering and transmission electron microscopy, immediately prior to performing assembly experiments.

We discovered, however, that incubating purified monomeric RSV CA (300 - 700 μM) at physiological or near-physiological temperatures (34-42 °C) over a period of several days promoted the formation of CA tubes. No tubes were identified in control samples incubated at 4 °C for equivalent time periods. It has been previously shown that both acidification ^19,27,51^ or the introduction of polyatomic anions like phosphate ^52^, promotes *in vitro* assembly of RSV CA into structures that mimic the authentic viral capsid. Linkage of RSV CA assembly to temperature has not been previously recognized. Because of its obvious biological relevance, we explored this temperature-dependent assembly in more detail. In our initial experiments RSV CA solutions (600-700 μM) were dialyzed for 16-24 hours against neutral pH buffers containing 25 - 1500 mM NaCl, at a temperature of 37 °C. To detect CA assembly, the dialyzed solutions were negatively stained, and examined by transmission electron microscopy (Fig 2).

**Fig 2.**
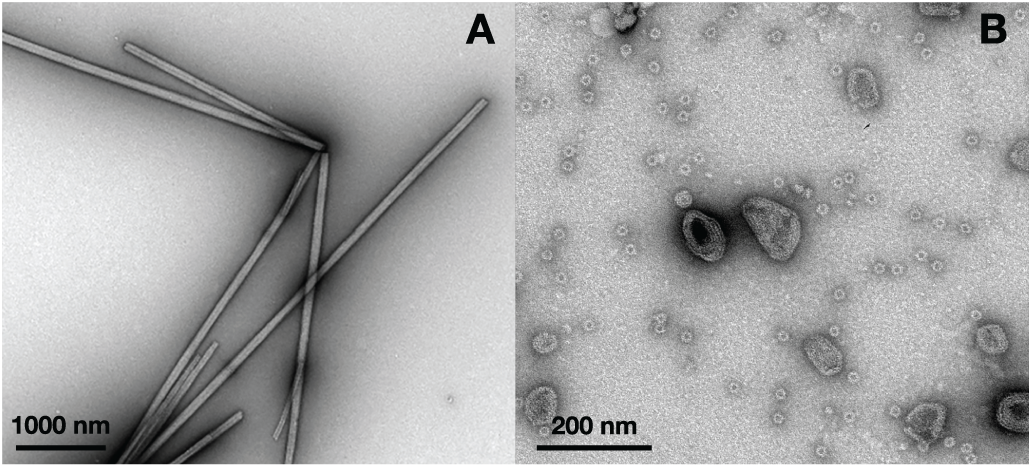
*In vitro* assembly of RSV CA at near-physiological temperatures. To promote assembly, purified RSV CA (600-700 μM in standard neutral pH storage buffer) was dialyzed against the same buffer, with variation of the NaCl concentration (25 - 1500 mM). Dialysis was carried out at a fixed temperature of 37 °C. The figure shows transmission electron micrographs of typical assembly products, negatively stained with uranyl acetate. (A) Low salt concentrations ( 25-150 mM) promote the exclusive formation of CA tubes. (B) Elevated salt concentrations (1500 mM NaCl) promote formation of CLPs and icosahedral particles.

Elevated temperature and low salt concentrations (25 −150 mM) led to the exclusive formation of CA tubes (Fig. 2A). These tubes appeared completely analogous to the well-characterized HIV-1 CA hexamer tubes which are readily assembled *in vitro* at high salt concentrations ^12,17,34,53–62^. Image analysis of tubes that had partially “unrolled” and were laid flat on the carbon support film showed convincingly that the tubes were built from capsid hexamers (Fig. S1). Tube formation appeared to be most efficient at sub-physiological salt concentrations. As the salt concentration was raised above physiological levels, RSV CA still assembled, but the outcome of the assembly process shifted. At 800 mM NaCl, a mixture of tubes and Core-Like-Particles (CLPs) was observed, while at 1500 mM NaCl tubes were no longer observed however CLPs and small spherical particles were abundant (Fig2B). The CLPs resemble authentic viral cores, which have a mean width of around 80 nm ^63^ and are highly polymorphic - some having a faceted and polygonal appearance, and others exhibiting extensive and continuous surface curvature ^63–65^. Based on the fullerene model of the mature retroviral capsid ^11^ the CLPs almost certainly contain both CA hexamers ^18,20^ and pentamers ^18,19^, while it is established that the small spherical particles are T=1 icosahedra, constructed from pentamers alone ^18,19^

Two firm conclusions can be drawn from these observational experiments. Firstly, assembly of RSV CA is strongly promoted at physiological temperatures. Secondly, the particular outcome of the assembly process is linked to the neutral salt concentration in the medium, which can apparently shift the proportion of hexamers and pentamers that are formed.

### Development of a Turbidimetric assay for tube assembly and disassembly

At physiological pH, temperature and salt concentrations RSV CA assembles exclusively into hexamer tubes. As these CA hexamers constitute the major building block of the orthoretroviral capsid we investigated the effects of temperature, protein and salt concentration on the tube assembly process.

To follow tube assembly in a more quantitative fashion we used a turbidimetric assay, in which the attenuation of incident light due to scattering by the tubes was measured using a conventional absorption spectrophotometer. Similar assays have been used to study HIV-1 CA tube formation ^56,58,62,66–69^ as well as the phosphate-driven assembly of RSV CA into CLPs ^50,52^. For long, thin tubes (relative to the wavelength of the incident light) the turbidity is a direct measure of the mass of protein that has polymerized ^70,71^. To increase both the rate and extent of tube formation an inert crowding agent (Ficoll 400) was introduced at the initiation of the experiment, by mixing stock solutions of crowding agent and protein (cf ^58^). Transmission electron microscopy was used to verify that tubes were the sole assembly product under all studied conditions.

### RSV CA tube assembly is strongly dependent on temperature, protein concentration and ionic strength

Initially, temperature was varied ( 6 - 34 °C), while the protein concentration (300 μM) and the NaCl concentration (75 mM) were held constant (Fig. 3A). Tube assembly was undetectable at low temperatures (≤13 °C) and barely discernible at room temperature (20 °C), even in the presence of the inert crowding agent. However assembly efficiency increased dramatically at higher temperatures (≥ 27 °C) in concordance with the earlier EM observations.

**Fig 3.**
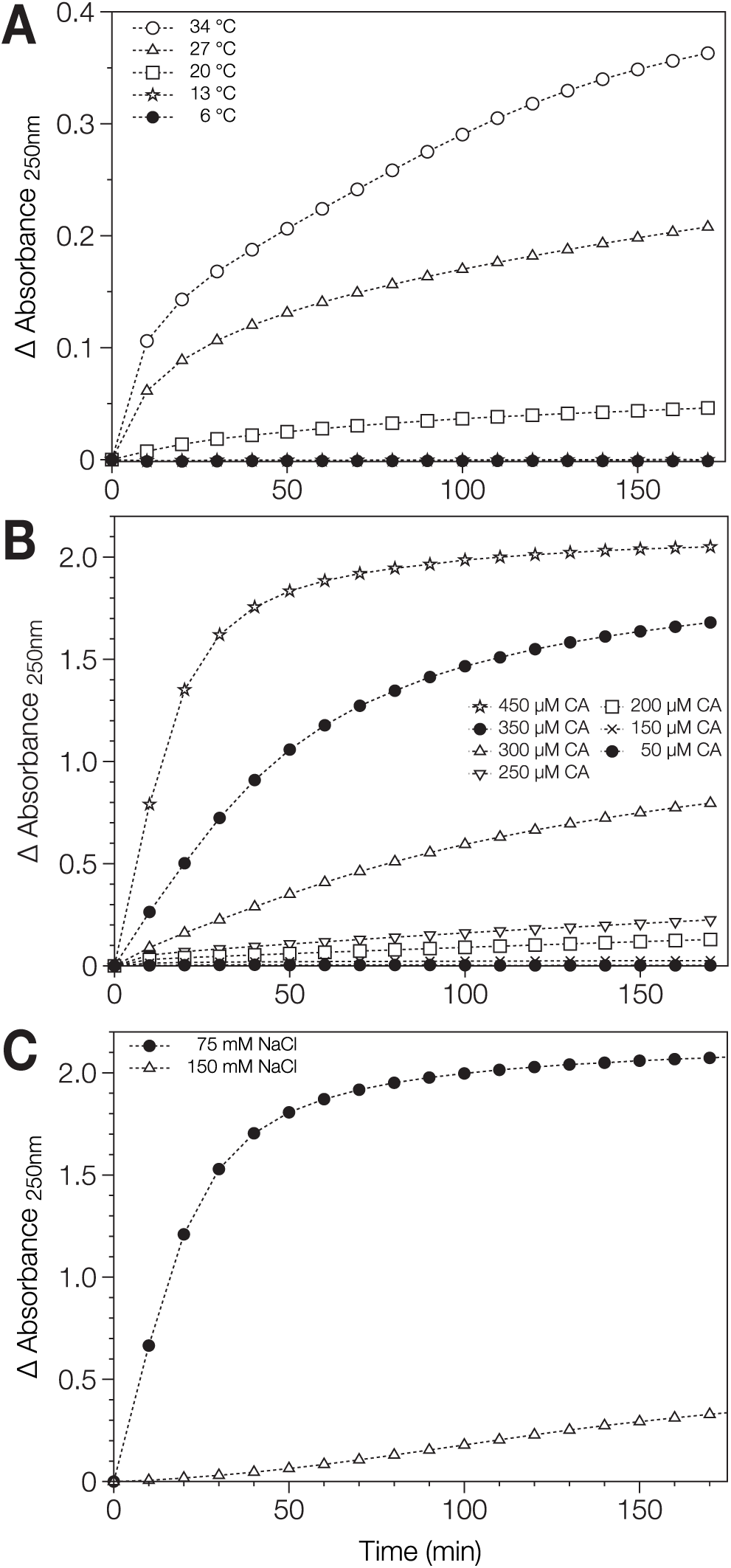
Assembly of RSV CA tubes followed using classical light scattering. (A) Dependence on Temperature. RSV CA (300 μM) in 12.5 mM MOPS/KOH pH 7.0, 75mM NaCl, 10%(w/v) Ficoll 400 was incubated at varying temperature. (B) Dependence on Protein Concentration. RSV CA (at varying concentration) in 12.5 mM MOPS/KOH pH 7.0, 75mM NaCl, 10%(w/v) Ficoll 400 was incubated at 34 °C. (C) Dependence on Neutral Salt Concentration. RSV CA (350 μM) in 12.5 mM MOPS/KOH pH 7.0, varying NaCl concentration, 10%(w/v) Ficoll 400 was incubated at 34 °C Absorbance at 250 nm was followed as a function of time in each case. The displayed absorbance data are the mean of replicate measurements made under each condition.

Subsequently, we examined the effects of protein concentration on tube assembly (Fig. 3B), working at a fixed temperature of 34 °C, where tube assembly is relatively efficient, and constant NaCl concentration (75 mM). Assembly was strongly linked to protein concentration, and there was clearly a threshold protein concentration below which assembly did not occur. The existence of a critical protein concentration suggests that tube formation is a nucleated process, in accord with simple theories of helical polymerization ^72,73^, and prior experimental ^19,52,59,74^ and theoretical ^75–77^ studies of orthoretroviral CA assembly.

Finally, we examined the effect of ionic strength on tube formation. RSV CA (350 μM) was incubated at a temperature of 34 °C, in the presence of sub-physiological (75mM) or physiological (150 mM) NaCl concentrations (Fig. 3C). In accord with the EM observations, tube assembly was much more efficient at sub-physiological salt concentrations suggesting that electrostatic interactions are central to tube formation. The effect of supra-physiological salt concentrations could not be meaningfully assessed using the light scattering assay, as tubes are not the exclusive assembly product under these conditions (Fig 2B). Collectively, the EM and light scattering results highlight the critical importance of charge screening effects in regulating RSV CA assembly, which was previously inferred from the ability of polyatomic anions like phosphate to promote *in vitro* assembly ^18,52^.

### RSV CA tubes assembled at high temperature only partially disassemble at low temperature

Light scattering was also used to explore the reversibility of the assembly process. A RSV CA solution was incubated for 3 hours under near-physiological conditions (34 °C, 75mM NaCl, pH 7.0), which facilitate tube formation. During this time the apparent absorbance steadily increased (Fig. 4), The temperature was then dropped to 13 °C, creating conditions that are not permissive for *de novo* tube assembly. Following the temperature drop, apparent absorbance decreased monotonically and plateaued at a value significantly greater than the initial value (Fig. 4). Hence temperature-dependent CA tube assembly is only partially reversible. The persistence of CA tubes at low temperatures, which do not permit their *de novo* assembly, shows that their formation remains thermodynamically favorable at these temperatures, and is only arrested by the large Gibbs energy barrier associated with nucleation.

**Fig 4.**
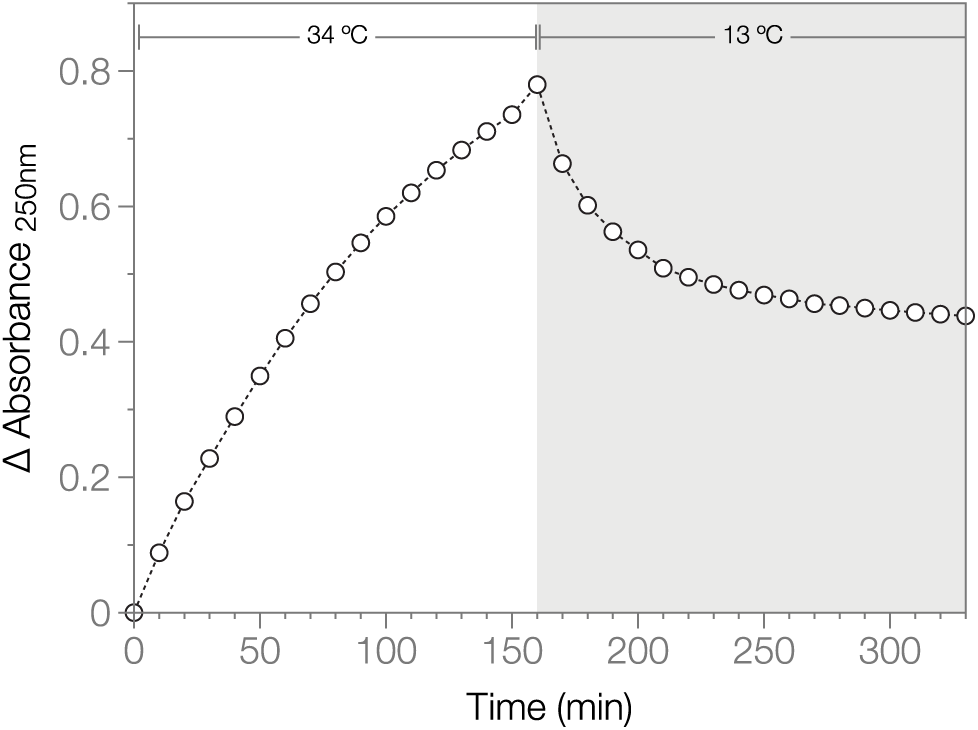
Disassembly of RSV CA tubes followed using classical light scattering. RSV CA (300 μM) in 12.5 mM MOPS/KOH pH 7.0, 75 mM NaCl, 10%(w/v) Ficoll 400 was incubated at 34 °C for 3 hours, while the apparent absorbance at 250 nm was monitored. The temperature was then dropped to 13 °C, and absorbance measurements continued for a further 3 hours. The displayed data are the mean of duplicate experiments.

### Mutagenic analysis confirms the critical importance of electrostatic interactions for RSV CA tube assembly

The sensitivity of RSV CA assembly to the neutral salt concentration suggests that electrostatic interactions are central to the assembly process. Such interactions are strongly modulated by the inclusion of mobile ions in the solvent. To further investigate this phenomenon, we manipulated the charge on the protein, by introducing pairs of charge-neutralizing point mutations at several locations on the surface of the NTD (Fig 5A). We then examined the *in vitro* assembly properties of the variant proteins. A total of three variants were constructed (K17A/R21A, R86A/R89A and R97A/R100A), with the paired mutations eliminating +2 elementary charges in each case. The K17A/R21A variant carries mutations in the first helix of CA, which helps form the central ring of both pentamer and hexamer (Fig 5A). The remaining two variants (R86A/R89A and R97A/R100A) carry mutations at the end of the 4th helix of CA, and in the flexible surface loop that follows. This loop - the structural analog of the cyclophilin A binding loop of HIV-1 CA - is located on the external face of the hexameric and pentameric rings that form the capsid, and is distal from the interfaces involved in forming the body of the ring (Fig 5A). As a consequence this region of CA is not in direct contact with other molecules in the assembled state.

**Fig 5.**
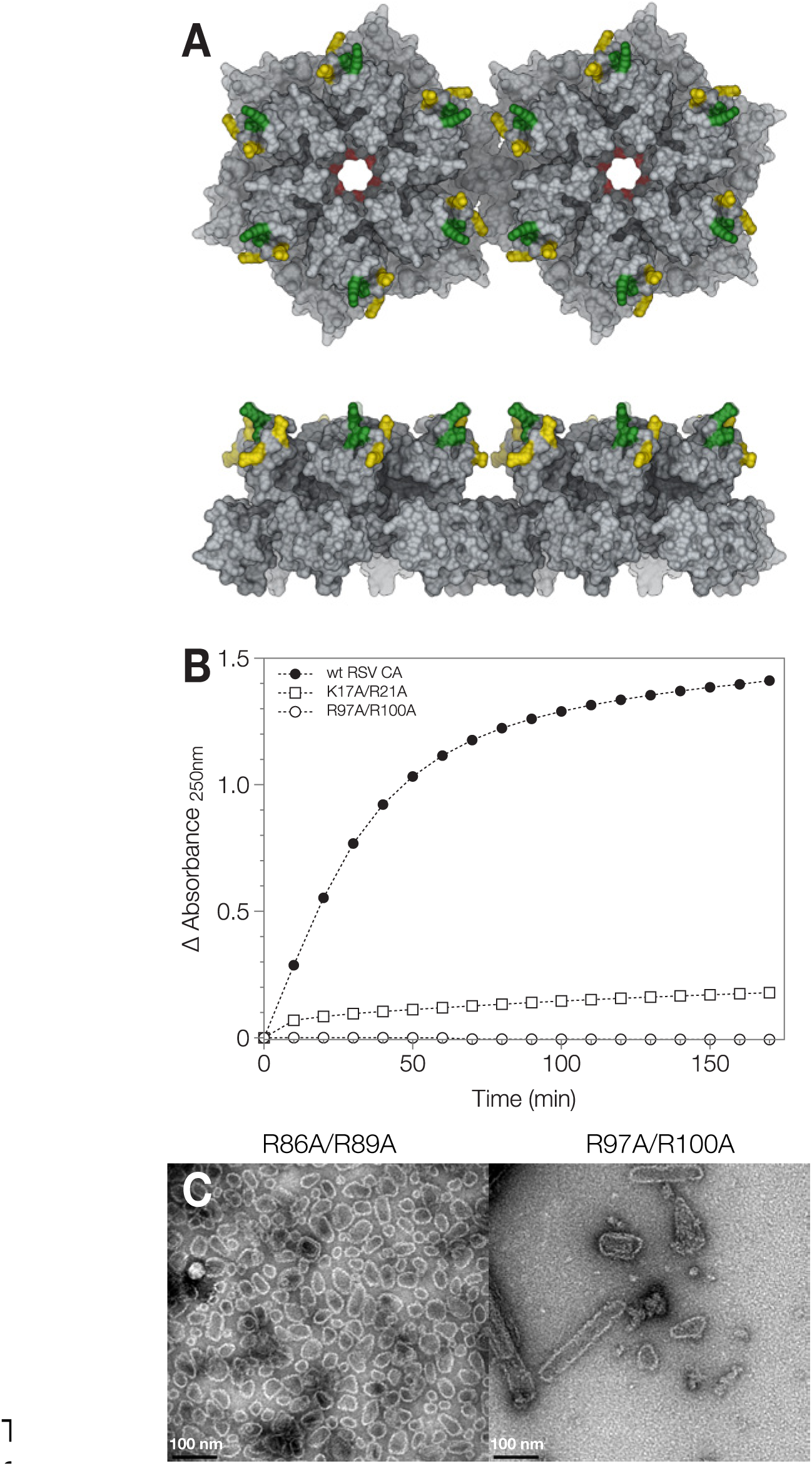
Effect of charge neutralizing mutations on RSV CA tube assembly. (A) The location of the charge neutralizing mutations introduced into RSV CA. Two RSV CA hexamers from a planar array (PDB ID: 3TIR) are shown in molecular surface representation, viewed from both the top and the side. Surface coloration indicates the location of mutations K17A/R21A (red), R86A/R89A (yellow) and R97A/R100A (green). (B) The mutations repress or abrogate tube assembly under physiological conditions. Wild type RSV CA and charge neutralized variants K17A/R21A and R97A/R100A (all 300 μM) were incubated in 12.5 mM MOPS/KOH pH 7.0, 75 mM NaCl, 10%(w/v) Ficoll 400 at 34 °C. Assembly was monitored using apparent absorbance at 250 nm. The displayed data are the mean of duplicate experiments. (C) The variants retain the ability to assemble into core-like-particles under non-physiological conditions. TEM images of typical assembly products are shown, following negative staining with uranyl acetate. To promote assembly, mutants R86A/R89A and R97A/R100A were dialyzed against neutral pH storage buffer at 37 °C, containing either 1.5M NaCl (R86A/R89A) or 0.8M NaCl (R97A/R100A). The program UCSF chimera ^110^ was used to generate parts of this figure, as well as Figs. 8 and 9.

We assessed the ability of the three variants to assemble into CA tubes by combination of electron microscopy and turbidimetric assay. The assembly of all three is severely impaired under near-physiological conditions. Both the variant and wild type proteins (600-700 μM) were dialyzed at 37 °C against pH 7 buffers containing 25-150 mM NaCl. After overnight incubation, wild-type CA tubes were abundant and readily visualized by electron microscopy. In contrast, no tubes were observed for variant K17A/R21A, and only occasional tubes observed for variants R86A/R89A and R97A/R100A, with no other assembly products apparent. This conclusion was reinforced by turbidimetric assay of variants K17A/R21A and R97A/R100A (Fig 5B) which shows that tube assembly is severely or completely abrogated, when compared with the wild type protein.

Despite this, the variants retained their ability to form capsid-like structures under non-physiological conditions that strongly promote assembly. For example, when dialyzed at 37 °C against pH 7 buffers containing very high salt concentrations (0.8 - 1.5M NaCl), all three variants formed CLP’s (Fig 5C), similar to the wild-type protein. Additionally, when transferred into acidic pH buffers, all three variants assembled into a mixture of sheets, CLPs and icosahedra (data not shown), similar to the wild type protein ^51^. Hence the impaired assembly of the variants under physiological conditions is not the trivial consequence of a major structural disruption, but almost certainly results from their altered electrostatic properties.

### The RSV CA tubes are multi-layered

To facilitate detailed structural analysis of the RSV CA tubes, we collected <u>Cryo-</u>EM images of the tubes suspended in a thin film of vitreous ice (Fig 6A). Examination of these images indicated that the tubes were invariably multi-layered, being composed of concentrically arranged layers of the capsid protein. Typical tubes had a diameter of ~100 nm and 3-4 concentric layers, but thicker tubes, with apparently as many as 7 or 8 layers, were sometimes observed. This phenomenon has considerable precedent. 2-D crystalline sheets of RSV CA hexamers assembled at acidic pH are always multi-layered ^20^. Multilayered CLPs also result from phosphate promoted-assembly of RSV CA ^18,52^. Additionally, HIV CA tubes assembled in the presence of elevated salt concentrations sometimes exhibit multi-layering (see e.g. ^12,59^).

**Fig 6.**
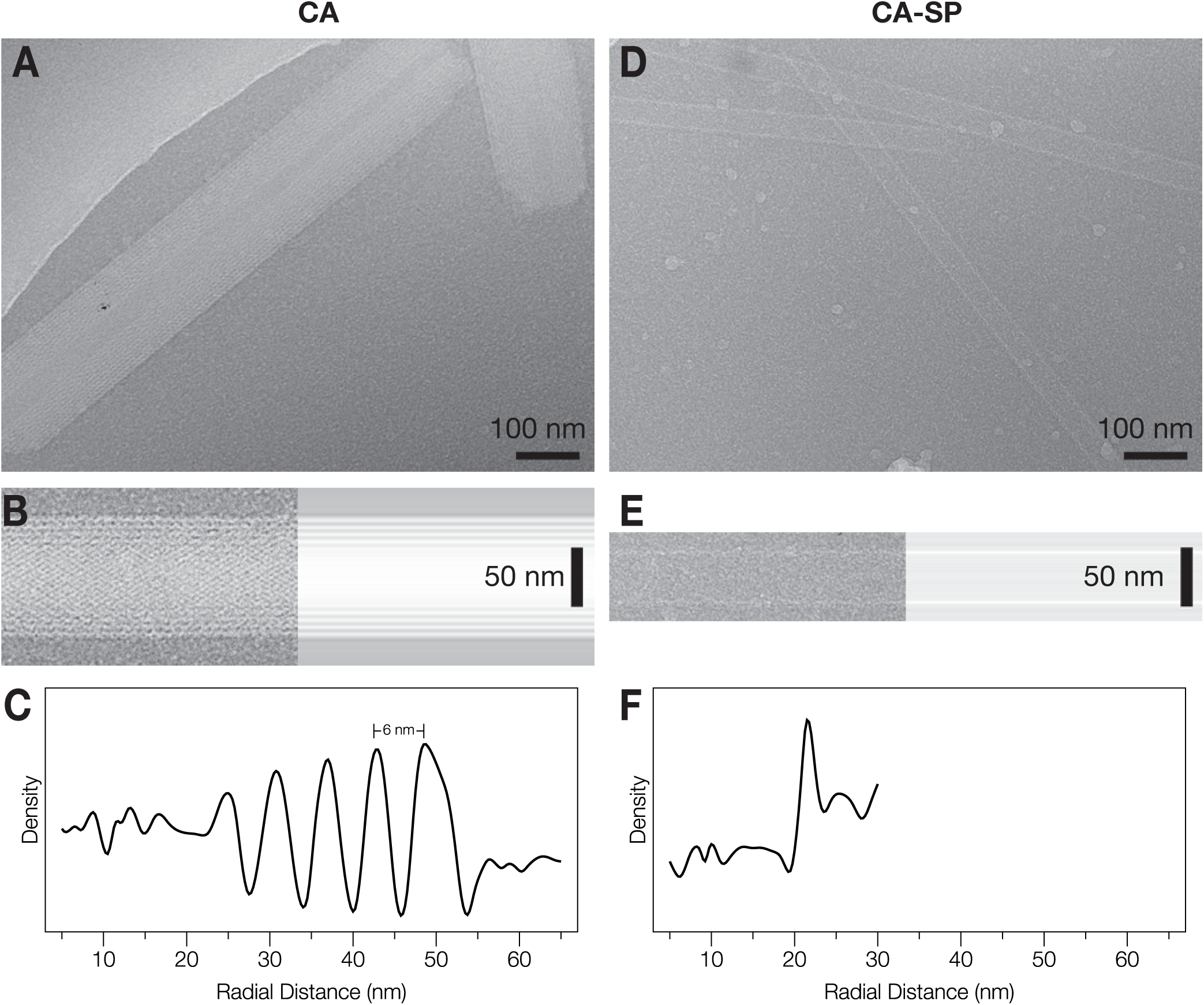
RSV CA and CA-SP Tubes imaged using Cryo-EM. (A) and (D) Images of tubes suspended in vitreous ice (B) and (E) Image of a tube (left), together with the longitudinally-averaged image of the same tube (right). (C) and (F) Mean radial density profiles for the tubes shown in (B) and (E) respectively. These were obtained from the longitudinally-averaged projection image via the inverse Abel transform. For this analysis, the images were not corrected for the effects of the contrast transfer function.

To help quantitate the layering phenomenon we calculated mean radial density profiles for individual tubes (Fig 6C). These can be readily estimated from projection images (Fig. 6B) by employing the inverse Abel transform (see materials and methods for details). The observed inter-layer spacing in the tubes (~6 nm, Fig. 6C)) is slightly larger than the 5 nm spacing between planar layers of RSV CA hexamers previously observed in 3D crystals ^20^.

### Addition of the spacer peptide to RSV CA blocks tube multi-layering

Multi-layering has been frequently observed when RSV or HIV CA are assembled *in vitro* ^12,18,20,52,59^ and multilayered (nested) capsids are also observed in virions ^64,78,79^. However, neither the physical cause nor the biological significance of multilayering is well understood. The RSV CA tubes assembled at high temperature are a good system for studying this phenomenon, as it is easy to detect and quantitate the multilayering effect.

Modifications at the C-terminus of RSV CA can influence the layering effects that are observed *in vitro* ^20,22^. For example, while full length RSV CA forms multi-layered hexamer sheets at low pH, a truncated variant lacking the disordered last 11 residues of the protein forms capsid mono-layers under the same conditions ^20^. For this reason, we investigated the effects of appending the C-terminal spacer peptide (SP) on RSV CA tube assembly. Because the retroviral protease cleaves the Gag polyprotein at several sites between CA and NC with differing rates, CA-SP (CA+12) is a persistent and detectable intermediate in the viral maturation process (Fig. 1), and is probably the major species involved in initiating mature capsid formation in the virion.

CA-SP assembles readily into tubes when incubated under near-physiological conditions. However, the CA-SP tubes were invariably single-layered (Fig. 6D, 6E, 6F), a result that is almost never observed when assembling CA alone. To confirm and extend this finding, we mixed CA and CA-SP in varying ratio and incubated the mixtures under identical conditions. We acquired cryo-EM images of the assembled material, and visually counted the number of single- and multi-layered tubes observed for each sample. Results are presented in Fig. 7 and Table 1. With inclusion of 10% CA-SP, a few single-layered tubes were observed (9%). However with inclusion of 30%-70% CA-SP, single-layered tubes became highly predominant (86-96%). Hence with respect to the assembly of RSV CA tubes under physiological conditions, the spacer peptide clearly acts as a layering determinant. Only a minor fraction of CA needs to be carrying the spacer peptide to effectively block multi-layered tube formation.

**Fig 7.**
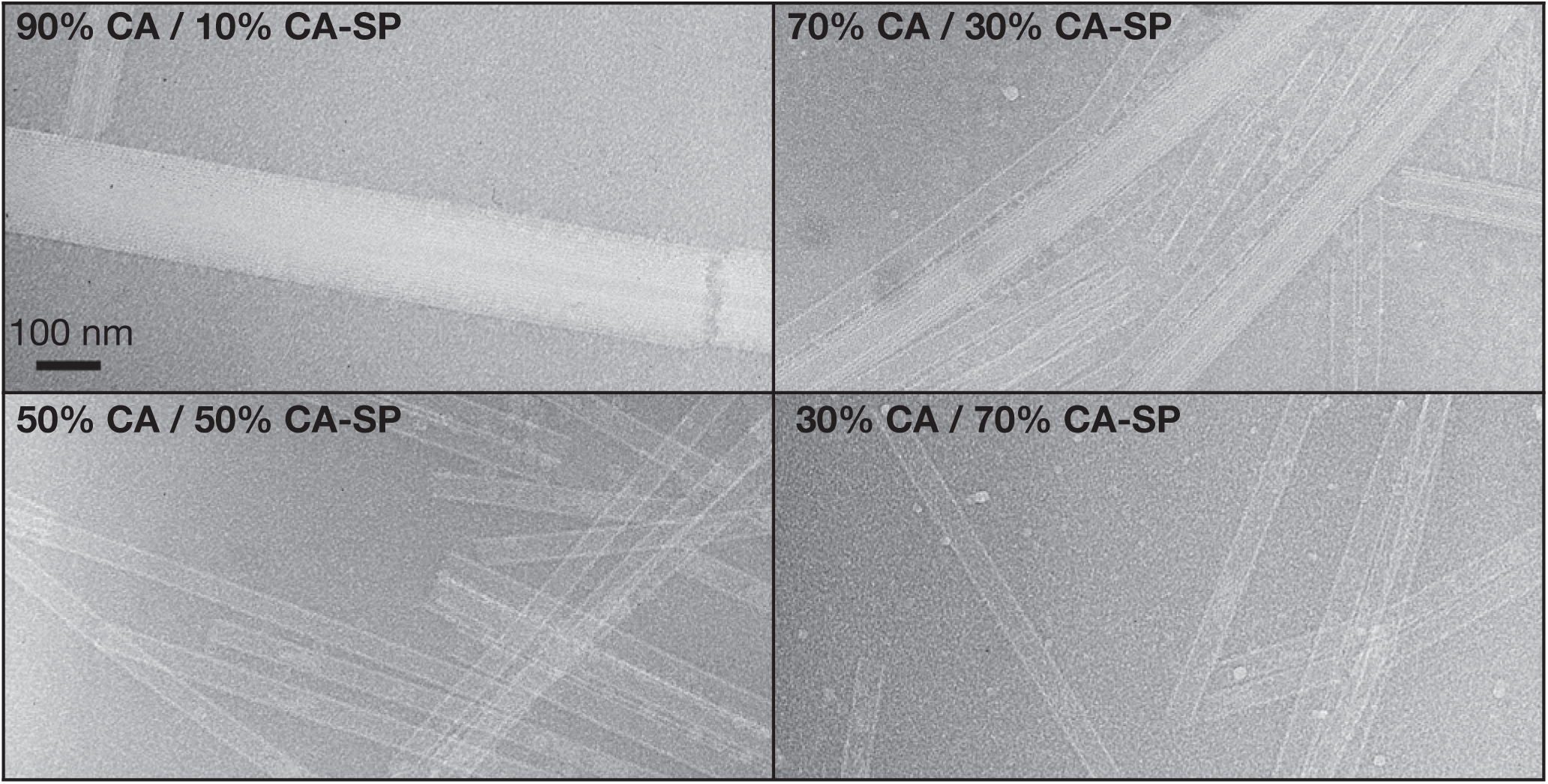
Cryo-EM images of tubes assembled from RSV CA / CA-SP mixtures. Images of typical tubes assembled from RSV CA / CA-SP mixtures at the indicated relative composition. To assemble the tubes CA/CA-SP mixtures (total protein concentration 570 μM in standard pH 7 storage buffer) were incubated at 37 °C for several days, before cryo-EM data collection.

**Table 1.** Tube Layering observed for CA-SP / CA mixtures.

| Composition | Number of Tubes Examined | % Single Layered | % Multi-layered |
| --- | --- | --- | --- |
| 10% CA-SP / 90% CA | 119 | 9 | 91 |
| 30% CA-SP / 70% CA | 166 | 86 | 14 |
| 50% CA-SP / 50% CA | 203 | 96 | 4 |
| 70% CA-SP / 30% CA | 243 | 93 | 7 |

If it is assumed that CA-SP and CA are recruited randomly into an assembling tube, then the statistics of incorporation into the hexameric building blocks of the tube will follow a binomial probability distribution. With 30% CA-SP, the probability that a single hexamer contains no CA-SP is 0.12, while the probability that two neighboring hexamers contain no CA-SP is only 0.01. The origins of the layering block could therefore be a structurally non-specific consequence of the relatively uniform distribution of SP over the inner face of the emerging hexamer lattice.

### 3D Image reconstruction of a single-layered RSV CA-SP tube

The generation of single-layered CA-SP tubes expedited analysis of the tube architecture using real space helical image reconstruction. To perform this analysis, we selected the tube image (Fig. 8A) whose Fourier transform (Fig. 8B) displayed layer lines to the highest resolution, and exhibited the lowest departures from mirror symmetry across the meridian. To determine the symmetry of the helix, we indexed the layer lines in the Fourier transform using standard procedures ^80–82^. However the diffraction pattern could not be unambiguously indexed in this fashion, and appeared consistent with 3 closely-related possibilities for the helical symmetry (See Materials and Methods for details, also Table S2, Fig. S2).

**Fig 8.**
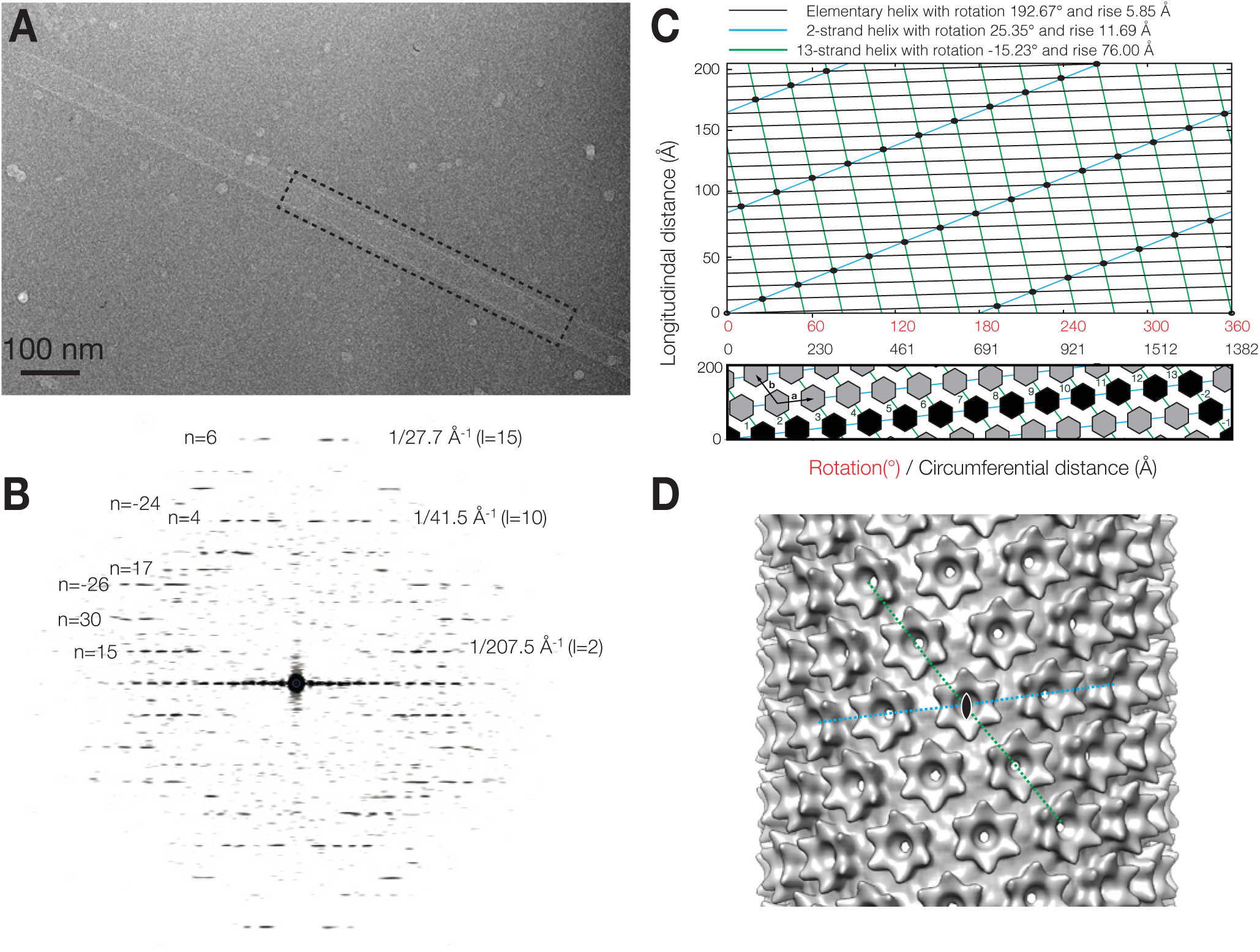
3D Image Reconstruction of a single-layered RSV CA-SP tube. (A) Cryo-EM image of the tube used for the 3D reconstruction (B) Power spectrum of the region outlined in (A), with some of the assigned Bessel orders (n) and layer lines (l) indicated. Clear layer lines are visible up to ~ 1/27.7 Å^-1^. (C) The surface lattice of the tube and the relationship between helical symmetry descriptions. The tube is created by a regular array of CA hexamers, arranged on a cylindrical surface. The diagrams represent radial projections of the structure, which corresponds to an unwrapping operation on the tube. The top panel illustrates the description of the helical lattice in terms of an elementary helix, a 2-strand helix, or a 13- strand helix. The scale for the vertical distance in this panel is expanded, for clarity. The bottom panel illustrates the description in terms of the generating vectors, **a** and **b**, of a near hexagonal 2D lattice, and the circumferential vector 13**a** - 2**b**. The black highlighting shows the components of the circumferential vector. The circumferential distance is calculated assuming a tube radius of 220 Å. (D) The final 3D reconstruction with local C6 symmetrization of the turrets applied. The view is down a 2-fold rotational symmetry axis, passing through the central hexamer. The disposition of the 2-strand and 13-strand helices shown in panel C are indicated. These coincide with the generating vectors of a near-hexagonal 2D lattice when the cylinder is unwrapped.

In order to discriminate among the 3 possibilities and refine the parameters of the correct helical screw rotation, we employed an iterative real space helical reconstruction scheme ^83^, as implemented in the program SPRING ^84^. In brief, after correcting for the effects of the contrast-transfer-function (CTF), the CA-SP tube image was divided into overlapping segments. These segments were aligned by projection matching, using a featureless cylinder as a reference. Then, making use of the helical screw rotation, the symmetrization procedure described by Sachse *et al* ^83^ was used to include all of the asymmetric units present in the tube image in the 3D reconstruction, with correct weight. The alignment and reconstruction procedures were then iterated until convergence. This scheme was followed for each of the 3 choices for the helical symmetry, in each case refining the helical rise and rotation by means of a grid search, while monitoring the consistency between the resulting reconstruction and the image.

Only one of the three possible helical symmetries generated a 3D reconstruction with surface features that were consistent with the known shape and dimensions of the retroviral CA hexamer. For this choice, the refined parameters of the elementary helix are a rise of 5.85 Å and a rotation of 192.67° (i.e. 1.868 units per turn), very close to those determined from initial Fourier analysis (Table S2). However it is probably more intuitive to describe the structure as a 2-strand helix, where each strand has a rise of 11.69 Å and a rotation of 25.35° (i.e. 14.20 units per turn), or as a 13-strand helix, where each strand has a rise of 76.00 Å and a rotation of −15.23° (i.e. 23.63 units per turn), with the individual strands related by the screw rotation of the elementary helix (Fig 8C, top). The significance of these particular helical families arises from the close relationship between the CA tubes, and the planar sheets of CA hexamers characterized previously ^20^. This relationship suggests an alternate description of the helical symmetry, involving the generating vectors of a planar lattice, and the circumferential vector, that describes the geometric relationship between a helical and planar lattice ^82,85,86^.

When a planar lattice is wrapped into a tube in a seamless fashion, the circumferential vector becomes the equator of the tube. As such, the circumferential vector represent the number of unit cells of the planar lattice required to complete a turn of the tube, and the concomitant displacement along the cell edge as this wrapping operation is performed. In this description, the helix is described by the circumferential vector 13**a** - 2**b**, where **a** and **b** are the generating vectors of the 2D lattice (the signs of the integer coefficients are based on helix polarity, according to convention ^82^) (Fig 8C, bottom). Assuming a tube diameter of 440 Å, the tube we have analyzed has the near-hexagonal unit cell dimensions a=98 Å, b= 96 Å, γ = 121°.

The hexagonal layer group symmetry (p6) which characterizes the planar CA sheets ^20^ is naturally lost as this structure is rolled into a tube. However inspection of the helical lattice, and the initial reconstruction, showed that 2-fold rotational symmetry axes that pass through the center of each hexagonal turret and the CTD dimer interfaces are maintained (i.e. the tubes are apolar structures possessing two fold rotational symmetry axes perpendicular to the principal helix axis).

To complete the 3D reconstruction (Fig. 8D), we enforced local 6-fold rotational symmetry around each hexagonal turret on the tube surface. In addition to improving the signal/noise ratio, this operation creates consistency with the known crystallographic structure of the turret ^20^, at the cost of introducing some perturbations into the reconstructed density on the internal surface of the tube. In this region, exact 6-fold symmetry cannot hold due to the variable displacement of the CTDs surrounding each turret. However as the floor of the tube is essentially smooth and featureless at the achieved resolution, this is inconsequential. The final reconstruction, included 784 hexamers and had an estimated resolution of 24 Å according to the FSC 0.143 criteria (Fig S3). The hand of the 3D reconstruction was inferred during fitting of the atomic model of the turret (Fig. 9, see discussion) though it cannot be reliably determined at this resolution.

**Fig 9.**
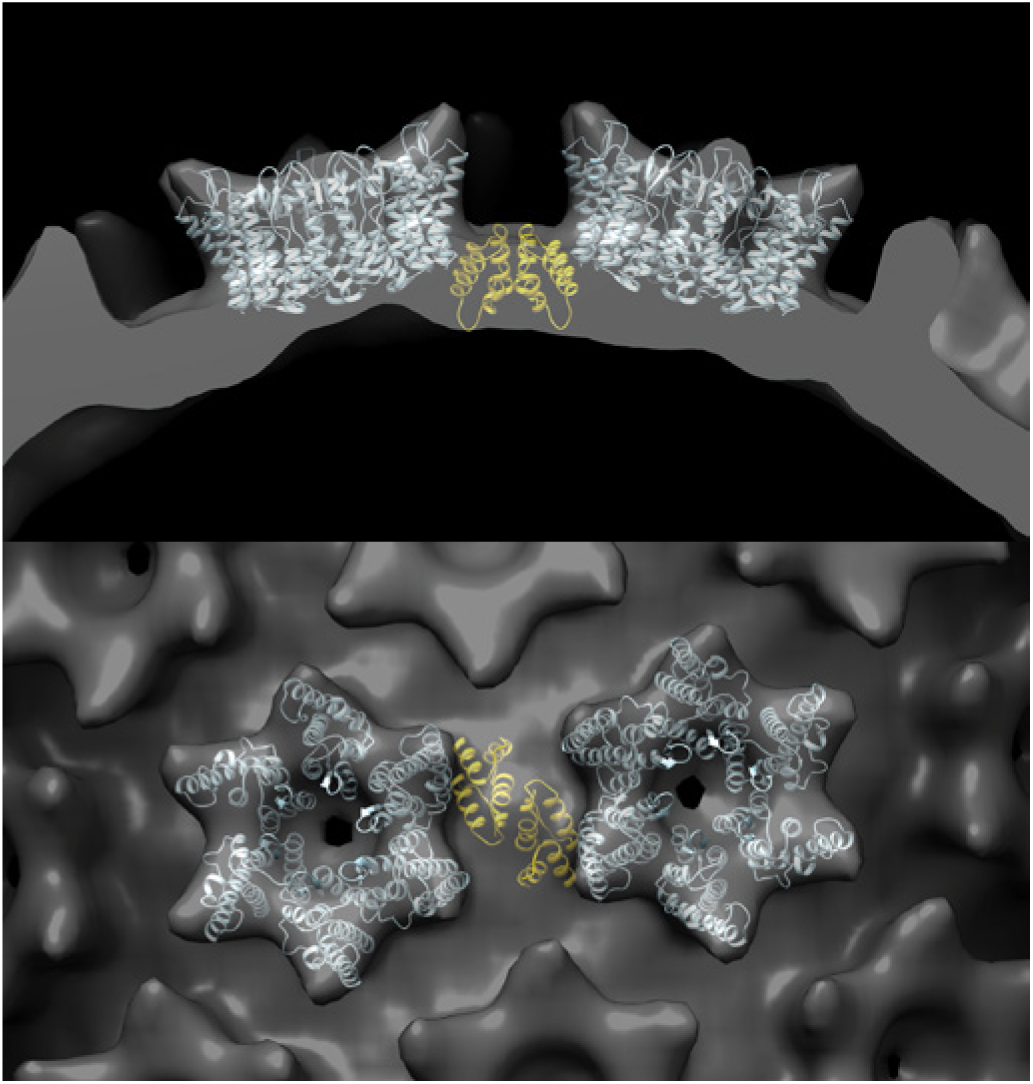
An illustrative molecular model, showing the connection between neighboring hexamers in the RSV CA-SP tube. NTD hexamers (light blue) were fitted into the 3D reconstruction using real space rigid body refinement with Phenix ^111^. A CTD dimer (Gold) was positioned underneath, in a position consistent with the the inferred interaction ^20^ between Ala 64 (NTD) and Asn 169 (CTD).The views displayed are parallel (top) and perpendicular (bottom) to the helix axis. Coordinates for the NTD hexamer and CTD dimer were taken from the pseudo-atomic model of the planar hexamer array (PDB ID: 3TIR).

## DISCUSSION

### Assembly of RSV CA hexamer tubes under physiological conditions

We have shown that incubation of RSV CA under physiological conditions is sufficient to promote assembly into tubes composed of CA hexamers, the major building block of the ortho-retroviral capsid (Fig S1, Fig. 8, Fig. 9). RSV CA tube assembly is a nucleated process (Fig. 3, Fig. 4), and the critical protein concentration required for assembly is relatively high. However, based on estimates of virion mass and volume ^63,87^, the CA concentration alone within viral particles would be in excess of 4mM, much higher than the critical concentrations required for nucleation. Hence the temperature-dependent assembly of RSV CA appears of direct biological relevance. In *vitro* assembly of RSV CA is also known to be promoted by both mild acidification ^19,27,51^ and the introduction of polyatomic anions like phosphate ^52^. However in neither case are hexamer tubes the exclusive result.

While RSV CA exists as a monomer under the conditions typical for *in vitro* analysis (neutral pH, ambient temperature, and physiological salt concentrations) ^19,27^, this state is metastable. At any appreciable protein concentration, capsid assembly is thermodynamically favored under these conditions, and is only held in check by the large Gibbs energy barrier associated with nucleation. We infer this from the equilibrium stability of RSV CA assembly products in physiological buffers, even when the original physical or chemical impetus for assembly is removed. For example, CA tubes generated at physiological temperature do not completely dissociate when the solution is shifted to lower temperature, and exist in equilibrium with unassembled protein (Fig. 4). These low temperatures do not support *de novo* tube assembly in the same buffer. Analogous observations were made for icosahedral CLPs that assemble readily at mildly acidic pH. These CLPs can be dialyzed into a neutral pH buffer without complete dissociation - conditions that are once again not permissive for *de novo* particle assembly ^19^. As a corollary, if suitable nucleating species are introduced into monomeric RSV CA solutions, assembly is immediately initiated ^19,52^.

Hence one of the essential aspects of the environmental perturbations now known to promote RSV CA assembly (acidification, introduction of polyatomic anions, and elevated temperature) is the lowering of this nucleation barrier. Only for acidification is there detailed insight into the nature of the nucleating species. In this case, protonation of a specific aspartic acid side chain (D191) on the surface of the CTD promotes CA dimerization, which is causally linked with higher order assembly ^51^. Enforcing CA dimerization, through introduction of an artificial disulphide bond across the dimer interface, creates a super-assembling RSV CA mutant ^51^. At present, the nature of the nucleating species being generated as the temperature is raised is unknown. Recent solution NMR studies of EIAV CA ^88^ have suggested that the CA conformation adopted within the hexameric ring is not significantly populated in solution, while studies on HIV CA have shown that the relative positioning of NTD and CTD is influenced by dimerization ^89^. Hence conformational shifting or selection could be a key step in the nucleation process ^88^. We note that the temperature-dependent enhancement of assembly may be unique to RSV CA as no comparable effect is apparent for HIV CA tube assembly ^54,59^. However Barklis and co-workers observed that brief incubation of HIV CA at elevated temperature accelerated tube formation at lower temperature, which they hypothesized was due to the creation of nucleating species ^59^.

Electrostatic interactions strongly influence RSV CA assembly at elevated temperature. The rate, extent and precise outcome of assembly are sensitive to charge screening by mobile solute ions. When the neutral salt concentration is sub-physiological, tube assembly is accelerated (Fig 3C). In contrast, when the neutral salt concentration is supra-physiological, tubes cease to be the sole assembly product, and CLPs ultimately predominate at high salt concentrations (Fig. 2B). The latter observation implies that the balance between hexamers and pentamers - the two fundamental building blocks of the capsid - is sensitive to the ionic environment in which assembly takes place. Supporting these observations we find that charge-neutralizing mutations on the surface of CA strongly modulate tube assembly. We eliminated positive surface charge by mutating pairs of residues both proximal (K17A/R21A) and distal (R86A/R89A and R97A/R100A) to the inter-molecular interfaces that form the hexamer array (Fig. 5A). These mutations repress or abrogate formation of CA hexamer tubes under physiological conditions (Fig. 5B), though the 3 variants still assemble under non-physiological conditions (Fig. 5C). The overall sensitivity of CA assembly to both mobile charge carriers in the solvent, as well as charged residues on the surface of the protein that are not in direct contact with other molecules in the assembled state, suggests that charge-charge interactions are very finely balanced to facilitate RSV capsid assembly *in situ*, consistent with prior observation for both RSV and HIV CA ^11,18^

Mutations of charged residues in first helix of CA, which helps form the central ring of both pentamer and hexamer, have previously been shown to influence retroviral capsid assembly. For example mutation R18A in HIV CA, appears to promote pentamer formation ^15,57^ and disfavour formation of hexamer tubes. Prevelige and coworkers ^67^ examined the effects of surface charge on HIV CA assembly, focussing on residues within the 2nd and 7th helices of CA located on the external face of the pentameric and hexameric rings, away from the interfaces involved in ring formation. Charge neutralizing and charge-reversing mutations in this region perturbed the assembly kinetics of HIV CA tube assembly *in vitro*. The variable electrostatic forces experienced by the sequence-diverse retroviral capsid proteins almost certainly contribute to the observed differences in the capsid assembly pathway, and the stability of the assembled capsid.

### Structural Analysis of RSV CA hexamer tubes

3D image reconstruction of a CA-SP tube (Fig. 8) shows that the RSV CA tubes formed at physiological temperatures are structurally analogous to the HIV CA hexamer tubes which are readily assembled at ambient temperature and supra-physiological salt concentrations ^12,17,34,53-62^. The hexameric turrets, which decorate the tube surface, result from relatively invariant NTD-NTD interactions. Each hexamer is connected to its immediate neighbours through CTD-CTD dimerization. Surface curvature can arise from the variable displacement of the six CTDs that surround each hexameric turret.

The mean diameter of single layered CA-SP tubes is ~ 45 nm, which we infer is associated with the most favoured curvature of the single-layered hexamer array. HIV CA tubes have similar mean diameters ^17^. However, in the absence of SP, the RSV CA hexamer tubes are invariably multi-layered (Fig. 6A & 6B), demonstrating the ability of the hexamer array to adopt varying mean curvatures. Continuously varying surface curvature is a feature of authentic viral capsids ^63-65^. Curvature is likely to originate primarily in simple pivoting movements at the NTD-CTD interface, possibly coupled with dislocations or displacements at the CTD-CTD dimer interface. This allows neighboring hexamers to tilt out of plane ^9 13,20,90^. Because of the 2-fold rotational symmetry associated with each hexamer in the tube, there are only three unique inter-hexamer connections. These must be associated with different NTD/CTD or CTD/CTD interfaces in order to create the necessary curvature. Crystallographic analysis of planar RSV CA hexamer arrays ^20^ suggests that Q169 (within helix α8 of the CTD) caps helix α4 of the NTD, creating a pivot that governs the relative displacement of NTD and CTD. A similar pivoting mechanism governing inter-domain motion may operate in HIV CA ^13^. It has not been established if dislocations at the CTD-CTD dimer interface also occur, or if each CTD-CTD dimer moves as a rigid body underneath the hexameric turrets it connects. Dimerization of the isolated RSV CA CTD is undetectable at neutral pH and sub mM CA concentrations ^51^ and some variation has been observed in structurally characterized RSV CA CTD dimers ^20,51^. This is also true for HIV CA ^9,12,44^ where CTD dimerization occurs with much higher affinity.

The limited resolution of the Cryo-EM reconstruction does not allow detailed modeling of the CTD displacements that allow the induction of curvature into the hexamer array. However we constructed an illustrative molecular model (Fig. 9) using the structures of the NTD hexamer and CTD dimer derived from prior crystallographic analysis ^20^. We fit the NTD hexamer structure into the density for the turret, and then positioned the CTD dimer underneath, such that Q169 from each CTD was in close proximity to Ala 64 in the correspondent NTD, thus maintaining the pivot on which inter-domain movement appears to be coordinated.

### The Spacer Peptide as a Layering Determinant

The spacer peptide acts as a layering determinant during hexamer tube formation. Under physiological conditions, CA forms multi-layered tubes, with multiple protein shells stacked concentrically upon one another (Fig. 6A, 6B, 6C). In contrast CA-SP forms only single-layered tubes (Fig. 6D, 6E, 6F). Mixture experiments, in which both CA and CA-SP are present in varying ratio, show that the longer species dominates the assembly phenotype (Fig. 7). Inclusion of CA-SP at even low amount blocks the formation of multilayered tubes (Table 1). This provides a relatively simple explanation for the frequent occurrence of multi-layered structures when CA is assembled *in vitro,* and the much rarer observation of these structures in authentic virions. In the virus, CA-SP is expected to be the predominant species when assembly is initiated (Fig 1).

It was previously concluded that RSV CA and CA-SP form essentially equivalent structures when assembled *in vitro* ^22^ - at least when using phosphate to promote CLP assembly. For example, in the presence of phosphate, both CA and CA-SP form multilayered icosahedral particles, having a T=3 external shell with a T=1 particle trapped inside ^22^. However for CA, the internal T=1 shell is in a fixed position with respect to the external T=3 shell, whereas for CA-SP it is not. Hence for *in vitro* assembly of icosahedral particles in phosphate buffers, the spacer peptide is clearly modulating the layering effect, while not abolishing it. This finding appears broadly consistent with our results, given the different methods adopted for promoting assembly, and the differing nature of the resulting assembly products. In addition, the presence of the spacer peptide significantly increased the rate of phosphate-promoted assembly into CLPs ^50^. Similarly, we observed that the spacer peptide accelerated tube assembly under physiological conditions (data not shown). The actual mechanism through which the spacer peptide exerts these effects remains unknown. The spacer peptide is quite hydrophobic in nature, and carries no ionizable side chains (Fig 1). Only a minor fraction of CA needs to be carrying the spacer peptide to effectively block multilayering (Fig. 7, Table 1) or accelerate capsid assembly ^50^. The acceleration of capsid assembly by the spacer peptide is also tolerant of conservative substitution (polyalanine mutation) ^50^. In the hexamer array, the spacer peptide must be localized around the trimer axis where the C-terminii of three CTDs are brought into close proximity. It seems probable that the spacer peptide is disordered in the hexamer tubes, consistent with prior observations on other structures that model the mature capsid ^22,49^. However the limited resolution of our 3D reconstruction leaves this question open. Plausibly, the acceleration of capsid assembly might arise from structurally non-specific hydrophobic interactions between spacer peptides, while the layering block may be steric in origin.

Irrespective of the mechanism, these observations suggest a model in which the transiently attached spacer peptide helps promote capsid assembly, and prevents formation of aberrant multilayered capsids. Presumably the spacer peptide can be more slowly trimmed from CA post-assembly, by the protease packaged inside the capsid shell. The enhanced rate of assembly of CA-SP ^50^ suggests that removal of the spacer peptide could have energetic consequences, effectively destabilizing the mature capsid and priming it for disassembly.

The *in vivo* relevance of the results awaits confirmation. It is noteworthy that while mutant viruses in which all or part of the spacer peptide is deleted appear to assemble and bud normally ^91,92^, such mutants are non-infectious. The lability of the capsid with respect to detergent treatment suggests that they have some irregularity in capsid organization ^91,92^. In the case of complete deletion, a condensed core was visualized in the mutant particles using thin section EM ^92^. At this point no high-resolution structural data is available which would help clarify the nature of the defect.

While the formation of multi-layered structures by self-assembling viral capsid proteins appears to be uncommon, it is not unique to the ortho-retroviruses. At suitable pH and ionic strength, the capsid protein of cowpea chlorotic mosaic virus (CCMV) forms multi-shelled capsids ^93,94^, likely instigated by the opposing charge carried by the inner and outer surfaces of the capsid shells. Mutated forms of the major bacteriophage T4 prohead component, gp22, result in formation of multi-layered proheads ^95^, and the protein will assemble into multi-layered tubes when other critical prohead components are inactivated or absent ^96^. Non viral proteins can also assemble in this fashion. For example muscle phosphorylase B can form doubled walled tubes when assembled *in vitro* ^97^.

### Summary

In summary, we show that incubation of RSV CA at physiological temperature, pH and neutral salt concentration results in the formation of tubes built from CA hexamers, the major component of the authentic capsid surface. Elevated temperature provides the trigger for CA assembly, at least in part by lowering the nucleation barrier, and hence the critical protein concentration that will support assembly. Using this biologically relevant assembly protocol, we show that the spacer peptide that is transiently appended to the C-terminus of CA functions as a layering determinant during formation of the capsid hexamer array. CA-SP, which is expected to be the predominant species when assembly of the mature capsid is initiated in the virion, forms only single-layered tubes, consistent with observations made on authentic viral capsids. In contrast CA alone forms multi-layered tubes, in which protein shells stack concentrically upon one another. Combined with earlier results, showing that the spacer peptide accelerates the rate of capsid assembly ^50^, the study demonstrates that the temporal control of Gag cleavage by the viral protease has potentially important consequences for capsid assembly and disassembly in the virus.

## METHODS

### Heterologous Protein Expression and Protein Purification

Recombinant proteins were produced by heterologous expression in *E. coli*., utilizing the IMPACT^TM^ expression system [NEB]. Expression constructs were created, encoding RSV CA and CA+12 (“CA-SP”) fused to a modified *Mycobacterium xenopi* GyrA intein. Following protein expression, DTT-mediated cleavage of the intein fusion proteins releases the naturally occurring CA protein variants (See Fig. 1, and associated discussion). Full details of the expression constructs are given in Supplementary Table S1. Plasmids encoding charge neutralizing RSV CA variants K17A/R21A, R86A/R89A and R97A/R100A were derived from the relevant parental plasmid using a two-stage PCR protocol ^98^, and verified by DNA sequencing.

Bacterial culture; lysis; affinity chromatography; and DTT-mediated cleavage of the intein fusion proteins was carried out as previously described ^51^. Working at 4°C, CA and variants were further purified by passing the cleaved protein (in 12.5 mM TAPS/KOH pH 9.0, 400 mM NaCl) back over a bed of chitin beads, to remove any residual intein molecules. In most cases the protein was then dialyzed for 8 hours against 25 mM sodium tetraborate/NaOH pH 10.0, 150 mM NaCl to fully hydrolyze the C-terminal thioester group resulting from intein cleavage. The elevated pH accelerates hydrolysis of the thioester ^99^. Finally the proteins were purified to homogeneity using size-exclusion chromatography (SEC) on Superdex 200 media [GE Healthcare] equilibrated in the standard protein storage buffer (12.5 mM MOPS/KOH pH 7.0, 150 mM NaCl, 0.5 mM NaN_3_, 0.25 mM TCEP.HCl). All the proteins eluted from the preparative SEC column at the volume expected of a monomer. Purified proteins were maintained at 4 °C until use. Protein samples that were subsequently concentrated using centrifugal devices (Vivaspin sample concentrators, GE Healthcare), were checked using dynamic light scattering and analytical SEC to verify that proteins did not assemble during the concentration step. Protein concentrations were estimated using UV absorption measurements at 280 nm ^100^. Dialysis of samples for preliminary EM analysis was carried out in 50-100 μL dialysis buttons (Hampton Research) employing a 3000 MWCO membrane.

### A turbidimetric assay for CA tube assembly

*In vitro* assembly of CA at elevated temperatures can be accelerated by the addition of a macromolecular crowding agent; Ficoll 400 (Sigma Aldrich). For the assembly assay, purified CA solutions (100-900 μM in standard storage buffer) were mixed with Ficoll 400 solutions (20 %(w/v) in a modified storage buffer containing 0-150 mM NaCl) at a volumetric ratio of 1:1, and a temperature of 4 °C. This resulted in final concentrations of 50-450 μM protein, 75-150 mM NaCl and 10%(w/v) Ficoll 400. The mixture was immediately transferred into a 1 mm path length quartz cuvette (Starna) and the apparent absorbance measured using a Cary 4000 UV-Visible spectrophotometer. The instrument was operated in dual beam mode with a matching Ficoll 400 solution in the reference cell. The samples were incubated at a fixed temperature for ~180 minutes, with a 240nm - 260nm wavelength scan recorded at ten minute intervals. Aliquots were removed from the cuvette for examination by negative-stain electron microscopy at the conclusion of the experiments.

### Electron Microscopy - Specimen Preparation and Data Acquisition

To prepare negatively stained specimens, 5 μL samples were applied to carbon-coated EM grids, rendered hydrophilic by glow discharge, and stained using 1-2%(w/v) uranyl acetate solutions. To prepare vitrified specimens, 5 μL samples were applied to perforated carbon EM grids (Quantifoil) that had been glow discharged in the presence of n-amylamine. Grids were blotted and plunged into a slush of liquid ethane using a semi-automated vitrification system (Vitrobot Mark IV, FEI) at 4 °C and 90 - 100 *%* relative humidity. All images were recorded using a Phillips Tecnai T12 microscope, equipped with a LaB6 filament, and operated at 120kV. Cryo-EM images of the tubes were recorded on SO-163 (Kodak) film at a nominal magnification of 42000x, at 1-2 μm underfocus. Selected images were digitized using a Nikon LS-9000 film scanner at a raster step size of 10.5 μM, corresponding to 2.5Å on the specimen.

### Electron Microscopy - Image Reconstruction

To compute longitudinally-averaged radial density profiles of the multi-layered tubes, it was necessary to accurately locate the tube axis in the projection images. The optimal axial position was identified by maximizing the correlation between the original image, and an image reflected through the cylinder axis. The image was then longitudinally averaged along the direction of the cylinder axis. From the longitudinally averaged projection data, the radial density profile of the cylinder was recovered by means of the inverse Abel transform, appropriate for reconstructing any object with a circularly symmetric cross-section. Inversion of Abel’s integral equation was done using the method of Deutsch and Beniaminy ^101^, which is stable in the presence of high levels of noise. Similar procedures have been used previously to recover the spherically-averaged radial density profiles of RSV virions and virus-like particles ^63,102^.

To analyze the Fourier Transform of the single layered CA-SP tube, a 700Å × 5358Å (280 × 2143 pixel) box encompassing the tube image was extracted using *e2helixboxer.py* in the EMAN2 package ^103^, after aligning the tube axis with the y-axis of the digitized image. This image was then floated and padded into a 10240 × 10240 Å (4096 × 4096 pixel) box to increase the sampling in Fourier space. The program *Bshow* within the Bsoft package ^104^ was used to inspect the Fourier transform and measure the positions of the visible layer lines and amplitude maxima. For each layer line, the module *segclasslayer* in SPRING ^84^ was used to measure and plot the phase differences across the meridian at the location of the first amplitude maximum. These measurements facilitate the determination of the parity (even or odd) of the Bessel order n. Bessel orders were estimated for a total of 15 well-defined layer lines, based on the distances from the meridian to the first amplitude maximum, and using an effective diameter of 440 Å for the tube. This estimate was approximate, due to the undulating nature of the tube boundary, the presence of defocus fringes, and the finite width of the tube. Additionally the parity of n was not apparent from the phase differences measured across the meridian. There were significant departures from the expected values of 0 or 180° for many of the layer lines, most likely due to the presence of out-of-plane tilt ( see ^105^) and/or non-uniform flattening of the tube. Indeed, based on the subsequent 3D image reconstruction, the out-of-plane tilt for the helix was estimated *a posteriori* as ~ 3°. As a result, unambiguous assignment of the Bessel order for most of the layer lines proved difficult.

The estimated values of *n* were plotted against the layer line number *l* (Fig S2). Because of the close correspondence between the *(n,l)* plot and the reciprocal lattice of the helix ^106^ the helical symmetry can be inferred directly from this plot. Given the uncertainties in n, we found that the *(n, l)* plot could be visually explained on the basis of 3 possible helical symmetries (Fig S2). Each possibility could be considered to arise from a 2-strand helix, with the strands having a pitch ~166 Å, but differing slightly in their geometry and relative disposition (Table S2). In one case (13.2 units per turn, Table S2) the two strands are related to each other by 2-fold rotational symmetry around the helix axis so that the subunits form a two-start helix with helical rotation/rise pair of 27.3°/12.5Å. In contrast, in the other 2 cases, (12.2 or 14.2 subunits per turn, Table S2) the 2-fold symmetry does not exist, and, instead, the subunits on both strands are part of the same elementary helix described by a helical rotation/rise pairings of 194.75°/6.8Å or 192.68°/5.85Å, respectively.

Iterative real space helical reconstruction was used to identify the correct symmetry and refine the helical parameters. For each possibility, reconstructions were performed while systematically varying the helical rise and rotation using the module *segrefine3dgrid* in SPRING ^84^ (Fig S3). After correcting for the effect of contrast-transfer-function (CTF) based on the CTF parameters estimated by CTFFIND3 ^107^, the image of the helix was partitioned into overlapping segments. The length of the segments was 1000 Å and the interval between segments 80 Å. Helical parameter refinement was done in two stages. The first stage involved searching a grid of ±0.5Å (interval 0.05Å) and ±2° (interval 0.2°), centered around the initial rise and rotation. The second stage employed a finer grid of ±0.2Å (interval 0.01Å) and ± 0.2° (interval 0.01°). In the *segrefine3dgrid* procedure, a full 3-D reconstruction was performed for each imposed helical symmetry, starting with a featureless hollow tube, and the resulting reconstructions were evaluated by monitoring agreement between the image data and the reconstructed volume (e.g. the cross-correlation between the projections of the reconstructed volume and the matching segments of the image).

Based on this analysis, scheme II could be discarded, while schemes I and III were indistinguishable by the criteria examined. This result may reflect some of the inherent ambiguities that exist in helical reconstruction ^108,109^. However inspection of the resulting 3D reconstructions showed that only scheme III resulted in a structure consistent with the known CA hexamer architecture.

To locally average the final reconstruction about the turrets on the tube surface, the optimal location of the 6-fold rotational symmetry axis was determined by a real space procedure. Sub-volumes were extracted, centered at each possible position within the angular and translational space defined by the helical repeat. For each sub-volume, a C6 symmetrized copy was created and correlated with the original sub-volume. From the resulting plot of cross-correlation versus centroid position, the location of symmetry axis was determined (Fig S3), and the local C6 symmetry imposed on the reconstruction.

## ACKNOWLEDGEMENTS

JKH and SAJ were supported by University of Auckland doctoral scholarships. We acknowledge the contribution of NeSI high-performance computing facilities to the results of this research. NZ’s national facilities are provided by the NZ eScience Infrastructure and funded jointly by NeSI’s collaborator institutions and through the Ministry of Business, Innovation & Employment’s Research Infrastructure programme. URL https://www.nesi.org.nz.

## AUTHOR CONTRIBUTIONS STATEMENT

SAJ performed the assays for tube assembly and the mutagenic analysis (Figs. 3–5). GDB prototyped the light scattering experiments and prepared samples for Cryo-EM. AD performed the 3D helical reconstruction (Figs. 8–9). JH collected the cryo-EM data (Figs. 6-7) and discovered the temperature dependence of RSV CA assembly (Fig. 2). AKM planned the experiments and assisted with writing the manuscript. RLK planned the experiments, performed some of the EM analysis (Fig. 7), and wrote the manuscript. All authors reviewed the manuscript.

## ADDITIONAL INFORMATION

The authors declare no competing financial interests.

